# Quantifying biomarker ambiguity using metabolic network analysis

**DOI:** 10.64898/2026.01.29.702649

**Authors:** Maggie Hinkston, Alexander S. Bradley

## Abstract

Molecular biomarkers preserved in rocks provide evidence about ancient life but interpreting them requires inference through multiple stages of information loss arising from phylogenetic, biosynthetic, and diagenetic ambiguity. However, biomarker specificity is typically assessed qualitatively rather than quantitatively. Here we formalize biosynthetic ambiguity as entropy over metabolic networks.

We introduce three metrics that quantify pathway-level information content: retrobiosynthetic complexity (ψ), normalized branch depth (λ), and fraction shared (σ). Analysis of 9,140 MetaCyc metabolites defines a three-dimensional specificity space for biomarker evaluation. Only 13% of multi-pathway compounds exhibited low complexity, distal divergence, and high pathway consensus. Lipid biomarkers span this specificity space heterogeneously: hopanoids cluster near the high-specificity region while sterols occupy intermediate territory. Diagnostic quality and lipophilicity are approximately independent, so the constraint on molecular paleontology is the limited chemical diversity among preservable compound classes rather than their biosynthetic properties.

This framework supports probabilistic biomarker interpretation by explicitly incorporating biosynthetic, phylogenetic, and diagenetic constraints.

## INTRODUCTION

Molecular fossils derived from lipids have been used to reconstruct life and environments across the span of Earth history^1,2^. Hydrocarbons retain structural and isotopic information inherited from their biological precursors, allowing inference about specific organisms or metabolic processes in ancient environments^3^. The emphasis on lipid-derived molecules reflects the exceptional preservation of hydrocarbons: although lipids encode less information than nucleic acids or proteins, they persist far longer in the sedimentary record^4^. For this reason, lipid-derived hydrocarbons provide an important window into ancient microbial ecosystems^5^, and similar molecules may be biosignatures in the search for life beyond Earth^6^. Interpreting these molecular fossils requires establishing connections between preserved hydrocarbons and their biological sources^5,7^ – relationships initially established through empirical lipid extraction^8–12^ and now increasingly predicted from genomic analysis of biosynthetic pathways^13,14^.

Despite these advances, biomarker interpretation still faces a persistent challenge: many lipids can be produced by multiple, phylogenetically distant organisms. This pattern arises through convergent evolution or horizontal gene transfer^1^, and the phylogenetic distribution of organisms capable of producing a specific molecule may change over the course of evolutionary history. The 2-methylhopanoids illustrate this challenge. Initially detected in cyanobacteria and methylotrophs^15^, and later in rocks and petroleum^16^, these molecules were widely adopted as biomarkers for cyanobacteria and for oxygenic photosynthesis^17^. Subsequent surveys showed that these molecules could be produced by a broader range of bacteria^18^, and that the genetic capacity for 2-methylhopanoid production is distributed across multiple lineages^14,19^.

This challenge is general: when encountering any fossil hydrocarbon, how can we determine which type of organism produced it? A better framing of this question may be: how confidently can we attribute a source to a biomarker? The degree of confidence depends on the specificity and distribution of the biosynthetic pathway. Molecules produced through a single, taxonomically restricted pathway provide more reliable source information than those that can be produced through multiple, widely distributed routes. Biomarker signals may reflect the evolutionary state of biosynthetic pathways themselves, not simply organismal presence or abundance^20^.

Framed this way, biomarker interpretation is fundamentally an inference problem. Given an observation (a molecule preserved in rock), we attempt to draw conclusions about the hidden state (the ancient organism, metabolism, environment, and diagenetic changes) that produced it. This inference must traverse multiple stages of information loss (Figure 1; Box 1): phylogenetic ambiguity arises when multiple organisms express the same pathway; biosynthetic ambiguity arises when multiple pathways produce the same molecule; and diagenetic ambiguity arises when multiple biomolecules yield the same preserved compound. Information theory provides tools for quantifying how much an observation reduces uncertainty about an unknown source^21^. Biomarker interpretation has usually relied on qualitative assessments of specificity or diagnostic value rather than quantitative measures of information content.

**Figure 1.**
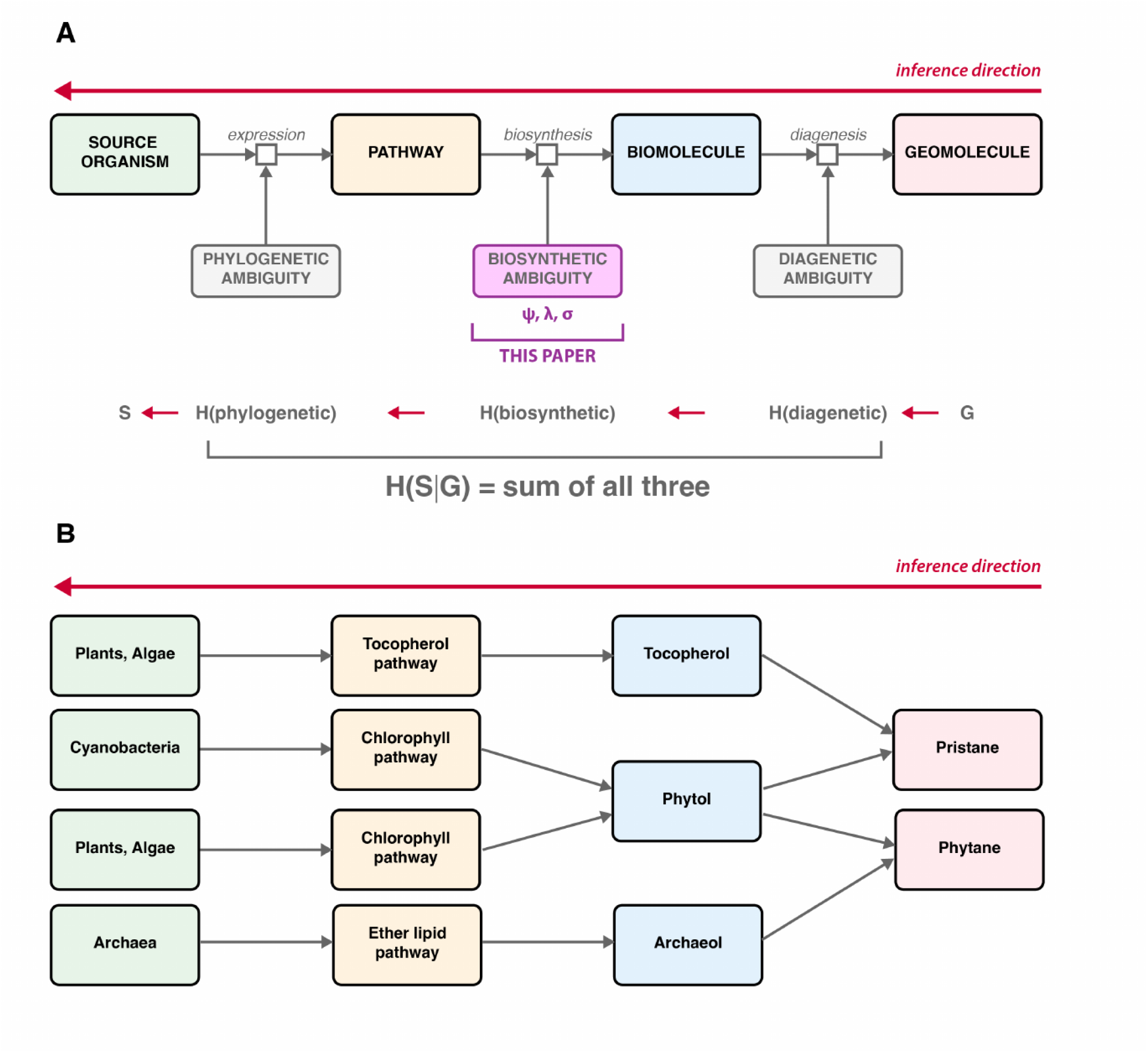
Biomarker interpretation as an inference problem. (A) Interpreting a preserved geomolecule requires reasoning backward through three stages of information loss. Source organisms express biosynthetic pathways, which produce biomolecules, which undergo diagenesis to yield preserved geomolecules. At each stage, many-to-one mappings introduce ambiguity: phylogenetic ambiguity (multiple organisms express the same pathway), biosynthetic ambiguity (multiple pathways produce the same molecule), and diagenetic ambiguity (multiple biomolecules yield the same preserved compound). The metrics introduced here (ψ, λ, σ) quantify biosynthetic ambiguity; the other two sources require separate treatment. (B) Pristane and phytane illustrate how ambiguity accumulates across stages. Phytol, produced via the chlorophyll pathway in cyanobacteria and algae, can yield either pristane or phytane depending on diagenetic conditions. Tocopherols (from plants and algae) provide an independent route to pristane, while archaeol (from archaea via the ether lipid pathway) provides an independent route to phytane. Observing pristane or phytane in ancient sediments thus leaves substantial uncertainty about both the precursor biomolecule and the source organism.

We present a quantitative framework for addressing biosynthetic ambiguity, based on three complementary metrics. Retrobiosynthetic complexity (*ψ*) quantifies the diversity of biosynthetic routes that could produce a molecule. Normalized branch depth (*λ*) quantifies where those alternative routes diverge, distinguishing proximal from distal pathway divergence relative to the target compound. Fraction shared (*σ*) quantifies the extent to which alternative routes traverse common biosynthetic intermediates, distinguishing compounds with conserved pathway backbones from those whose alternatives proceed through largely independent sequences of reactions. By analyzing the structure of metabolic networks in reverse, from product to precursors, we systematically evaluate which biomarkers provide the strongest constraints on their biological sources.

The metrics *ψ, λ*, and *σ* capture independent aspects of biosynthetic organization and together define a three-dimensional specificity space. Ideal diagnostic compounds exhibit low pathway diversity, distal divergence, and extensive shared biosynthesis. A biomarker’s interpretive value depends on how strongly its presence constrains possible origins: a molecule produced by a single pathway provides far stronger constraints on source attribution than one accessible through many alternative routes. Quantifying biosynthetic specificity enables Bayesian approaches to biomarker interpretation, in which pathway-based priors can be updated with contextual evidence such as depositional environment, isotopic signatures, or co-occurring molecules, to yield probabilistic source assessments rather than deterministic assignments.

This study tests three related hypotheses: (i) that biosynthetic ambiguity can be quantified as entropy over metabolic networks; (ii) that pathway diversity, branching position, and pathway overlap encode independent constraints on biomarker interpretability; and (iii) that compounds with optimal biosynthetic specificity are rare and systematically biased toward poor geological preservation.

### Box 1 | Biomarker interpretation as an inference problem

Interpreting molecular fossils requires reasoning backward from observation to origin: given a preserved compound, what can we infer about the organism that produced it? This inference must traverse multiple stages of information loss (Figure 1). Source organisms express biosynthetic pathways, which produce biomolecules, which undergo diagenesis to yield the geomolecules we observe. At each stage, many-to-one mappings introduce ambiguity that limits what can be recovered.

**Three sources of ambiguity.** Phylogenetic ambiguity arises when multiple organisms express the same biosynthetic pathway. For example, the chlorophyll pathway operates in cyanobacteria, algae, and land plants. Biosynthetic ambiguity arises when multiple pathways produce the same biomolecule; cholesterol can be synthesized through dozens of post-lanosterol routes. Diagenetic ambiguity arises when multiple biomolecules yield the same preserved compound: pristane derives from both phytol (via chlorophyll) and tocopherols, while phytane can derive from phytol or archaeol. Each many-to-one mapping constrains what can be inferred about earlier stages.

**Quantifying uncertainty with entropy.** Information theory^22^ provides tools for measuring these constraints. The entropy H(X) of a random variable quantifies uncertainty about its value^21^. For biosynthetic pathways, under a uniform prior:

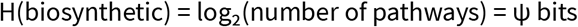

In Figure 1A, total uncertainty in source attribution H(S|G) is partitioned into phylogenetic, biosynthetic, and diagenetic components. The metrics introduced in this study quantify only the biosynthetic term H(biosynthetic): ψ measures its magnitude, while λ and σ describe how that entropy is distributed along the pathway graph. Retrobiosynthetic complexity (ψ) directly measures entropy over biosynthetic routes. A compound with ψ = 0 bits has a single pathway (no biosynthetic ambiguity), while ψ = 10 bits corresponds to ∼1,000 routes. ψ quantifies biosynthetic ambiguity only and represents one component of total source uncertainty.

**Mutual information and diagnostic value.** Mutual information I(X; Y) quantifies how much observing Y reduces uncertainty about X. For biomarkers, I(source; molecule) represents diagnostic value – how strongly a compound’s presence constrains inference about its origin. An ideal biomarker would have I(source; molecule) = H(source), meaning the observation uniquely determines the source. A non-diagnostic compound has I(source; molecule) ≈ 0. Diagnostic value depends on all three ambiguity sources. Low ψ (few pathways) is necessary but not sufficient: if those pathways are phylogenetically widespread or if the biomolecule converges diagenetically with other precursors, information about the source is still lost. The metrics introduced here (ψ, λ, σ) quantify biosynthetic ambiguity; phylogenetic and diagenetic ambiguity require separate treatment.

**Partitioning biosynthetic information.** The normalized branch depth metric (λ) indicates where alternative pathways diverge: high λ means pathways share most steps and diverge only near the product; low λ means early divergence near primary metabolism. The fraction shared (σ) quantifies overlap among routes: high σ indicates a conserved biosynthetic backbone; low σ indicates largely independent pathways.

These metrics define a diagnostic profile:

| Metric | Relation to H(biosynthetic) |
| --- | --- |
| $\psi$ (complexity) | Total biosynthetic entropy |
| $\lambda$ (norm. branch depth) | Spatial localization of entropy within pathways |
| $\sigma$ (fraction shared) | Partitioning between shared vs. distinguishing |

Hopanoids such as diploptene (ψ = 2, λ = 0.50, σ = 0.86) exemplify favorable profiles: few pathways sharing extensive structure. Cholesterol (ψ = 166, λ = 0.44, σ = 0.07) shows how similar λ can mask different architectures - its many pathways proceed through largely independent intermediates, providing weak biosynthetic constraint despite similar branching depth.

**Implications.** Low biosynthetic ambiguity is neither necessary nor sufficient for high diagnostic value - phylogenetic distribution is ultimately determinative. A compound produced by many pathways all restricted to one lineage can be highly diagnostic, while a compound produced by a single widespread pathway cannot. However, biosynthetic specificity establishes the complexity of the phylogenetic mapping problem: low ψ means fewer pathways must be surveyed across taxa, making assessment tractable, while high ψ compounds require characterizing the distribution of many independent routes. Quantifying biosynthetic ambiguity is thus a necessary first step toward comprehensive biomarker evaluation.

## RESULTS

We developed three complementary metrics to quantify biosynthetic ambiguity (Figure 2). Retrobiosynthetic complexity (ψ) measures pathway diversity – the entropy, in bits, over alternative biosynthetic routes to a compound. Normalized branch depth (λ) measures where those routes diverge, ranging from 0 (proximal divergence near the target compound) to 1 (distal divergence near terminal metabolites). Fraction shared (σ) measures the extent to which alternative routes traverse common intermediates. These metrics define a three-dimensional specificity space for biomarker evaluation.

**Figure 2.**
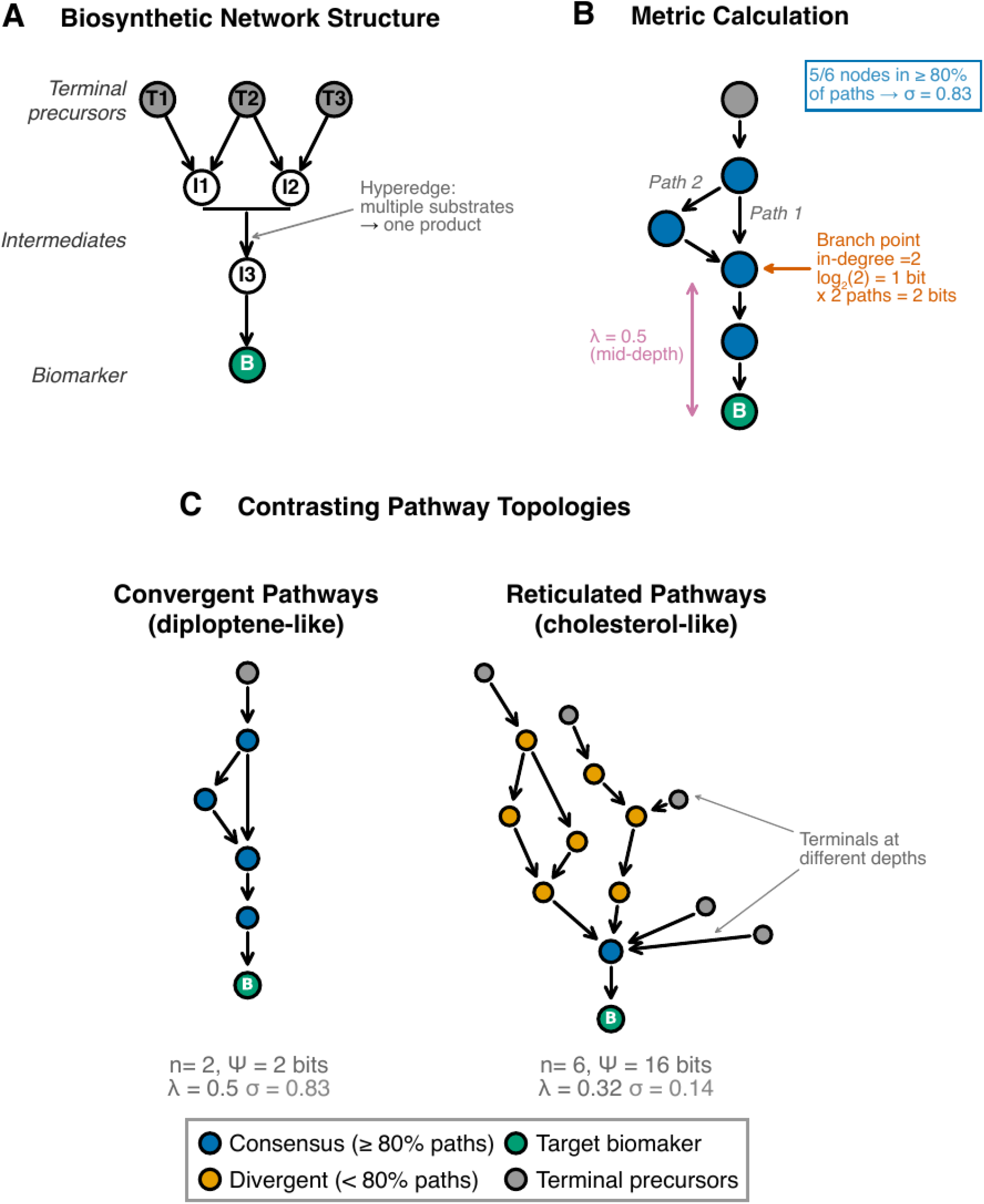
Quantifying biosynthetic ambiguity using metabolic network metrics. (A) Biosynthetic networks are represented as directed hypergraphs, where nodes represent metabolites and hyperedges represent reactions that may have multiple substrates converging to produce a single product. Terminal metabolites (gray) are biosynthetic starting points; intermediates (white) connect terminals to the target biomarker (green). (B) Calculation of the three metrics illustrated on a convergent pathway. Retrobiosynthetic complexity (*ψ*) accumulates at branch points where multiple reactions can produce a metabolite; here the single branch point (in-degree = 2) contributes log_2_(2) = 1 bit, yielding *ψ* = 2 bits across both paths. Normalized branch depth (*λ*) measures the normalized depth at which branching occurs (*λ* = 0.5 indicates mid-pathway branching). Fraction shared (*σ*) quantifies the proportion of nodes appearing in ≥80% of paths; here 5 of 6 nodes are consensus nodes, yielding *σ* = 0.83. (C) Contrasting pathway topologies. Convergent pathways (left) have few paths that share extensive biosynthetic machinery (high *σ*), characteristic of compounds like hopanoids. Reticulated networks (right) have multiple entry points at different depths, generating many paths through largely independent intermediates (low *σ*), characteristic of compounds like sterols. Metrics shown are calculated from the schematics as drawn: convergent pathway (n = 2 paths, *ψ* = 2 bits, *λ* = 0.50, *σ* = 0.83); reticulated network (n = 6 paths, *ψ* = 16 bits, *λ* = 0.32, *σ* = 0.14).

### Global Metabolic Network Analysis

The global small molecule metabolism network constructed from MetaCyc comprised 10,467 unique metabolites connected by 32,193 reactions. The network displays scale-free structure typical of biological metabolic networks, with most compounds participating in few reactions while a minority function as highly connected hubs. After excluding 602 compounds lacking identifiable biosynthetic pathways and 725 compounds with more than 10,000 biosynthetic paths, 9,140 compounds remained for analysis.

### Biosynthetic complexity is bimodally distributed

Among the 9,140 analyzable compounds, 4,510 (49.3%) had single biosynthetic pathways (ψ = 0 bits), while 4,630 (54.3%) had multiple pathways. The complexity distribution was bimodal (Figure 3): 2,198 multi-pathway compounds showed constrained biosynthesis (ψ ≤ 8 bits), while the remaining 2,432 (26.6% of all compounds) formed a high-complexity group (ψ > 8 bits, median = 70 bits, 80% range = 12– 9,101 bits) spanning nearly three orders of magnitude.

**Figure 3.**
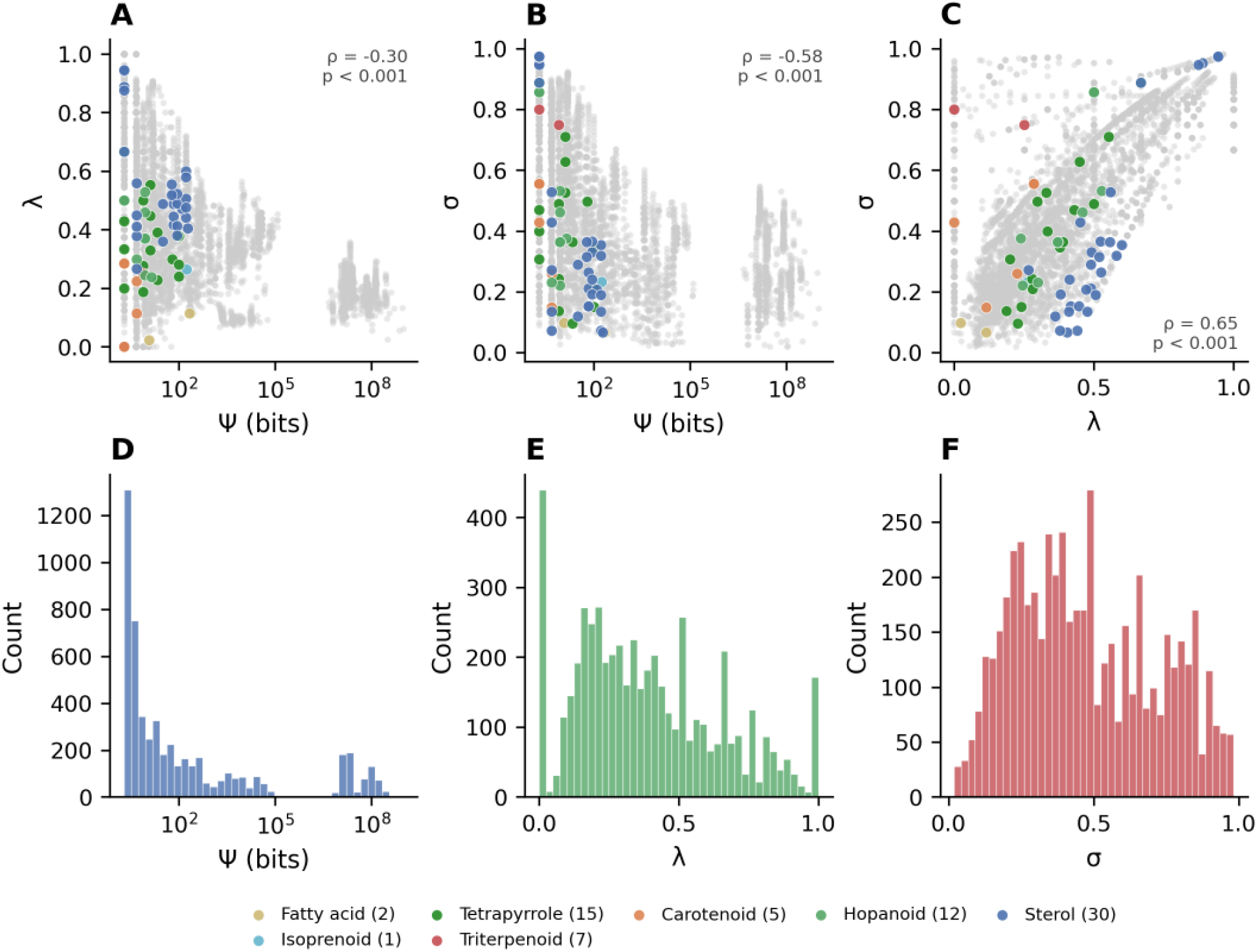
Distribution of biosynthetic metrics across the MetaCyc metabolome. (A–C) Pairwise relationships among retrobiosynthetic complexity (*ψ*), normalized branch depth (*λ*), and fraction shared (*σ*) for 4,630 multi-pathway compounds. Grey points show all compounds; colored points indicate biomarker classes. Spearman correlation coefficients (ρ) are shown. (A) *ψ* vs *λ* shows weak negative correlation (ρ = −0.29), indicating pathway diversity and branching position capture largely independent aspects of biosynthetic organization. (B) *ψ* vs *σ* shows moderate negative correlation (ρ = −0.57), with high-complexity compounds tending toward lower pathway consensus. (C) *λ* vs *σ* shows positive correlation (ρ = 0.64), reflecting the geometric constraint that distally-branching pathways share more proximal steps. (D–F) Distributions of individual metrics. (D) *ψ* distribution is right-skewed (median 9 bits) with most compounds showing low complexity. (E) *λ* distribution spans the full range with median 0.40. (F) *σ* distribution shows median 0.50, with approximately half of compounds exhibiting high pathway consensus.

### Compounds with optimal diagnostic properties are rare

Among 4,630 compounds with multiple biosynthetic pathways, normalized branch depth (λ) ranged from 0 to 1 (median = 0.35), with 8.2% showing proximal branching (λ = 0) and 13.9% showing distal branching (λ ≥ 0.7). Fraction shared (σ) ranged from 0.02 to 0.98 (median = 0.46), with 44.9% of compounds showing high pathway consensus (σ > 0.5).

Combining all three metrics identifies compounds with optimal diagnostic properties: low ψ (< 32 bits), high λ (> 0.7), and high σ (> 0.5). Only 714 compounds (13.3% of multi-pathway metabolites) met all three criteria. These compounds have conserved, distally-branching architectures well suited for source attribution.

### ψ, λ, and σ are largely orthogonal

To characterize relationships among the three metrics, we analyzed pairwise correlations across all compounds with multiple pathways (n = 4,630). Retrobiosynthetic complexity showed weak negative correlation with normalized branch depth (ρ = -0.16, p < 0.001) and moderate negative correlation with fraction shared (ρ = -0.55, p < 0.001), indicating that pathway diversity and branching architecture are largely independent, while high-complexity compounds tend toward lower pathway consensus.

Normalized branch depth and fraction shared showed moderate positive correlation (r = 0.65, p < 0.001): compounds with distal branching naturally tend toward higher consensus because shared proximal steps constitute more of each route. However, the correlation is imperfect (r^2^ = 0.43), and σ provides information beyond λ alone.

Decile analysis confirmed these patterns. From lowest to highest ψ deciles, mean λ decreased from 0.42 to 0.19 and mean σ from 0.64 to 0.33. High-complexity compounds tend toward proximal branching and lower pathway consensus, but considerable variation exists within each complexity class; biomarker evaluation therefore benefits from all three metrics.

Retrobiosynthetic complexity is independent of structural complexity. To test whether retrobiosynthetic complexity recapitulates molecular structure, we compared ψ, λ, and σ against 13 structural descriptors capturing molecular size (heavy atom count, molecular weight), complexity (Böttcher score^23^, Bertz CT^24^) synthetic accessibility (SA Score^25^), lipophilicity/polarity (logP^26^, TPSA, H-bond donors/acceptors), stereochemistry (stereocenters, Fsp3^27^), and flexibility (ring count, rotatable bonds) for 7,380 compounds with valid structures (Supplementary Methods, Supplementary Figure 1). No structural descriptor correlated strongly with ψ (all |ρ| < 0.18; Supplementary Figure 1). A linear model using all 13 descriptors explained only 8% of log_2_(ψ) variance (adj. R^2^ = 0.077). The SA Score, which estimates synthetic accessibility from structural fragments24, showed no significant correlation with ψ (ρ = −0.019, p = 0.255). In contrast, λ showed moderate correlation with ring count (ρ = 0.345) and stereocenters (ρ = 0.262), consistent with the expectation that structurally complex molecules require deeper biosynthetic pathways. Retrobiosynthetic complexity thus captures pathway-level information - the multiplicity and topology of alternative routes – that cannot be inferred from molecular structure alone.

### Biomarker classes occupy distinct regions of ψ–λ–σ space

Sterols (n = 30) exhibited moderate complexity (mean ψ = 66.5 bits, range 2–192), moderate λ (mean λ = 0.55), and moderate fraction shared (mean σ = 0.38), consistent with multiple post-lanosterol pathways through divergent intermediate sequences. The C_30_ sterols (24-isopropylcholesterol and 24-n-propylcholesterol) show favorable λ and σ values (λ = 0.94, σ = 0.97), with branching only at the terminal alkylation step.

Hopanoids (n = 12) showed low complexity (mean ψ = 6.8 bits, range 2–14) with variable fraction shared (mean σ = 0.53, range 0.22–0.89). The simplest hopanoids – diploptene and diplopterol – exhibit high σ (0.86), reflecting the conserved pathway from squalene to the hopanoid skeleton. Extended hopanoids with side-chain modifications (e.g., bacteriohopanetetrol, σ = 0.22) show lower consensus as each modification step introduces pathway alternatives. Despite internal variation, hopanoids exhibit substantially lower complexity than sterols, consistent with their empirical utility as bacterial markers.

Fatty acids (n = 2) showed high complexity (mean ψ = 113 bits) with proximal branching (mean λ = 0.07) and low fraction shared (mean σ = 0.08), consistent with limited diagnostic utility. Carotenoids (n = 7) exhibited low complexity (mean ψ = 2.3 bits) with proximal branching (mean λ = 0.13). Tetrapyrroles (n = 15) showed moderate complexity (mean ψ = 31.9 bits, mean λ = 0.34).

Archaeal ether lipids (n = 11) represent uniquely constrained biosynthesis, with all compounds showing single pathways (ψ = 0 bits). This reflects the recently characterized tetraether synthase (Tes) pathway^28^. GDGTs are highly diagnostic as archaeal markers.

These patterns demonstrate that biosynthetic architecture varies systematically with compound class: hopanoids and GDGTs occupy the high-specificity region, sterols occupy intermediate space with considerable internal variation, and fatty acids show high complexity consistent with diverse biosynthetic origins.

### DISCUSSION

Chemical selection, not biosynthetic limitation, shapes the biomarker record. Lipid biomarkers are selected for geological preservation primarily by lipophilicity. Biomarkers in our dataset have dramatically higher logP (median 7.7) and lower polar surface area (median TPSA = 20 Å^2^) than the general metabolome (logP = 0.8, TPSA = 104 Å^2^), confirming the well-established chemical filter imposed by diagenesis. However, this filter does not systematically exclude compounds with favorable diagnostic properties. To test this, we defined a composite diagnostic quality score as the sum of z-scored metrics (z(−ψ) + z(λ) + z(σ)), where higher values indicate biosynthetically simpler, more convergent, and more shared pathways. Diagnostic quality correlates weakly and positively with lipophilicity across the metabolome (Spearman ρ = +0.10), and biomarker diagnostic quality is statistically indistinguishable from the metabolome as a whole (Mann-Whitney p = 0.75). Biosynthetic specificity and chemical persistence are approximately orthogonal: a compound’s lipophilicity provides almost no information about its retrobiosynthetic complexity, and vice versa.

The biomarker record therefore samples limited chemical diversity rather than systematically excluding diagnostic compounds. Molecular paleontology operates within constraints imposed by organic geochemistry – we interrogate the subset of biosynthetic diversity that survives, not the subset that would be most informative. Lipid biomarkers span the full range of diagnostic quality, from the 6th to the 100th percentile, but each compound class occupies a narrow region of ψ–λ–σ space (Figure 4; Table 1). Triterpenoids achieve the highest diagnostic quality among biomarker classes (93rd percentile), while hopanoids combine low retrobiosynthetic complexity (median ψ = 9 vs 65 for sterols; Mann-Whitney p = 0.02) with high pathway consensus. These class-level differences are biologically meaningful and reflect the evolutionary histories of specific pathway architectures rather than properties related to preservation chemistry.

**Table 1.** Summary statistics for biosynthetic metrics across biomarker classes. Retrobiosynthetic complexity (ψ), normalized branch depth (λ), and fraction shared (σ) for 86 biomarkers grouped by compound class. n, number of compounds analyzed per class.

| Class | n | Mean $\psi$ | Median $\psi$ | Min $\psi$ | Max $\psi$ | Mean $\lambda$ | Mean $\sigma$ |
| --- | --- | --- | --- | --- | --- | --- | --- |
| Sterol | 30 | 66.5 | 64.7 | 2 | 191.6 | 0.547 | 0.383 |
| Hopanoid | 12 | 6.8 | 9 | 2 | 14 | 0.435 | 0.534 |
| Triterpenoid | 7 | 2.9 | 2 | 2 | 8 | 0.488 | 0.852 |
| Carotenoid | 7 | 2.3 | 2 | 0 | 5 | 0.13 | 0.564 |
| Archaeal lipid | 11 | 0 | 0 | 0 | 0 | 0 | 1 |
| Tetrapyrrole | 15 | 31.9 | 13 | 2 | 101 | 0.338 | 0.372 |
| Fatty acid | 2 | 113.4 | 113.4 | 11.9 | 214.8 | 0.069 | 0.083 |
| Isoprenoid | 1 | 174.7 | 174.7 | 174.7 | 174.7 | 0.264 | 0.233 |
| Algal hydrocarbon | 1 | 0 | 0 | 0 | 0 | 0 | 1 |

**Figure 4.**
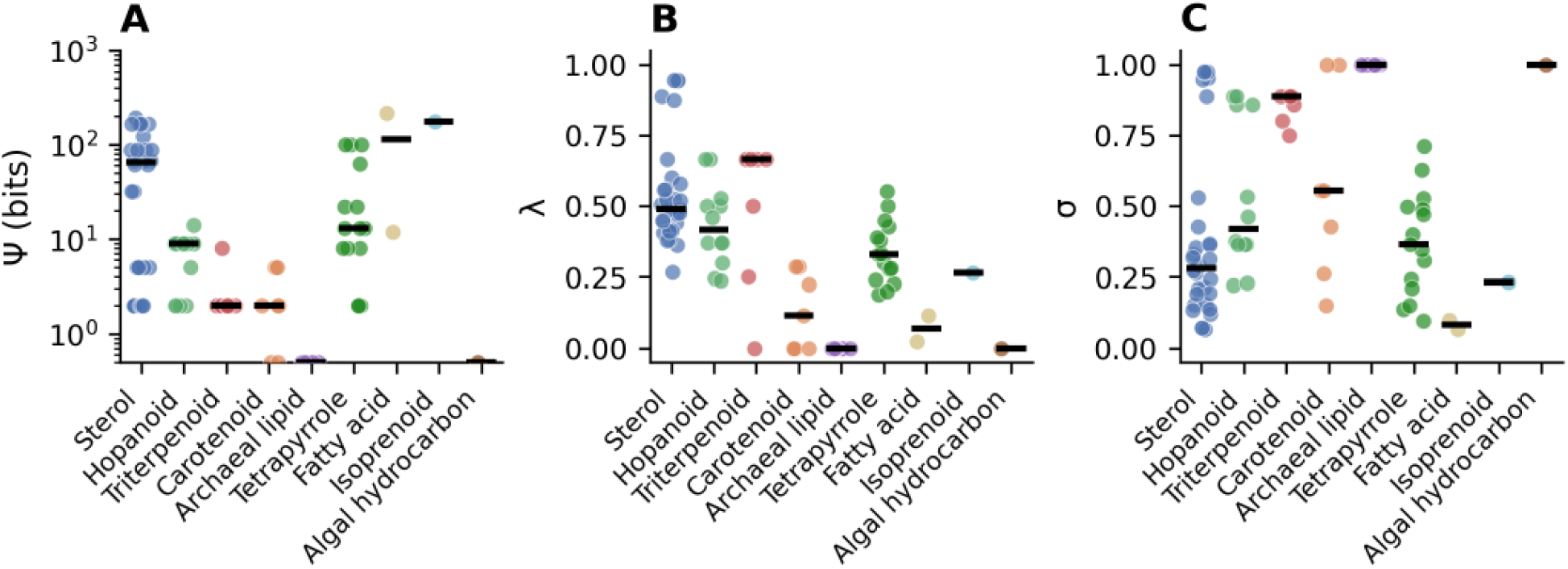
Biosynthetic metrics across biomarker classes. (A–C) Distribution of retrobiosynthetic complexity (ψ), normalized branch depth (λ), and fraction shared (σ) for 86 biomarkers across 9 compound classes. Each point represents an individual compound; horizontal bars indicate class medians. (A) Retrobiosynthetic complexity varies by several orders of magnitude across classes. GDGTs and isoprenoids show minimal pathway diversity (ψ ≤ 2 bits), while fatty acids exhibit high complexity (median ψ > 100 bits). (B) Normalized branch depth shows substantial within-class variation, with sterols and hopanoids spanning nearly the full range. (C) Fraction shared distinguishes compound classes with conserved biosynthetic backbones (triterpenoids, GDGTs, hopanoids; σ > 0.5) from those with largely independent pathways (fatty acids, porphyrins; σ < 0.2). Complete metrics for all biomarkers are provided in Supplementary Table S1.

This independence between biosynthetic informativeness and chemical persistence means that biomarkers with favorable diagnostic properties exist but are distributed unevenly across compound classes. Expanding the biomarker toolkit through novel compound classes or improved analytical sensitivity for trace polar lipids could access regions of ψ–λ–σ space not currently represented in the geological record and find utility in modern environments.

### The biomarker inference chain

Interpreting molecular fossils requires reasoning through sequential stages of information loss. A preserved geomolecule derives from one of potentially many precursor biomolecules, synthesized through one of potentially many pathways, by one of potentially many organisms. Working backward, the interpreter faces three compounding uncertainties: diagenetic ambiguity (many biomolecules yield the same preserved compound), biosynthetic ambiguity (many pathways produce the same biomolecule), and phylogenetic ambiguity (many organisms possess the same pathway). Each stage filters information; reliable source attribution requires that at least one stage provides strong constraint.

The metrics presented here quantify biosynthetic ambiguity - the middle term in this chain. Biosynthetic complexity determines whether phylogenetic mapping is tractable. For compounds with few pathways, surveying which organisms possess each route is feasible, while for compounds with many pathways, comprehensive surveys become impractical. Low ψ does not guarantee taxonomic restriction, but it makes the question answerable.

Pathway metrics can mislead when phylogenetic distribution does not track pathway structure. A biosynthetically constrained compound may nonetheless be produced by distantly related organisms through horizontal gene transfer or convergent evolution of the relevant enzymes. Hopanoids illustrate the distinction between biosynthetic and phylogenetic ambiguity. Despite low ψ (mean 6.8 bits) and high σ (0.86 for diploptene), hopanoid production spans bacteria, ferns^29^, and fungi^30^ - phylogenetically distant groups that share a conserved squalene-hopene cyclase. Low biosynthetic complexity indicates pathway surveys are tractable; phylogenetic distribution determines whether such surveys yield diagnostic specificity. Conversely, GDGTs demonstrate when pathway and phylogenetic constraint align: tetraether biosynthesis proceeds through a single known route that remains restricted to archaea; these compounds are therefore reliable domain-level markers.

Diagenetic ambiguity compounds the problem in the other direction. Steranes in ancient sediments derive from diverse sterol precursors through loss of functional groups and stereochemical rearrangement; hopanes similarly represent multiple hopanoid sources. The specificity of a geomolecule depends on its own biosynthetic complexity and on how many precursors funnel into it during diagenesis. A biosynthetically constrained compound that shares its diagenetic fate with many other precursors provides weaker evidence than one whose preservation pathway is also distinctive.

Quantifying biosynthetic ambiguity thus addresses one term in a multi-term problem (Box 1). It identifies which compounds warrant the substantial effort of phylogenetic and diagenetic analysis – and which are intrinsically ambiguous regardless of how thoroughly those analyses are pursued.

### Toward probabilistic biomarker interpretation

Retrobiosynthetic complexity allows Bayesian approaches to biomarker interpretation. Using Bayes’ theorem, the probability that a detected biomarker (B) derives from a particular source (S) is:

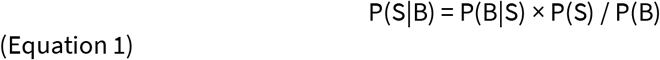

Our metrics quantify pathway-level ambiguity, providing one component of this calculation. Compounds with low ψ have constrained biosynthesis, making the likelihood terms P(M|S) tractable to estimate through phylogenetic surveys of the relevant enzymes. But the real utility comes from integrating multiple lines of evidence. Consider 2-methylhopanoids, now known to be produced by cyanobacteria, alphaproteobacteria, acidobacteria, and other lineages^14,19,31^. Detected alone, these compounds provide weak source constraint. But environmental context modifies the prior: in shallow-water carbonates where cyanobacteria thrive, P(cyanobacteria) increases substantially compared to deep-water or anoxic settings. Co-occurring biomarkers provide additional constraints through joint probability calculations – detecting 2-methylhopanoids alongside cyanobacterial carotenoid derivatives would multiplicatively increase confidence in a cyanobacterial source. Host rock lithology, depositional environment, and stratigraphic context all contribute information that can be formally integrated.

Under this approach, confidence in source attribution can be updated rather than collapsed when new producers are identified. In a categorical framework, the demonstration that unicellular rhizarians can synthesize 24-isopropylcholesterol^32^ justifiably negates C_30_ steranes as diagnostic sponge biomarkers. Under a probabilistic Bayesian interpretation, the relevant question is not whether alternative producers exist, but how pathway structure, together with geological context and other evidence, modifies the posterior probability of source attribution. The C_30_ sterol case demonstrates how our approach reframes taxonomic debates. Rather than asking whether 24-isopropylcholesterol is ‘diagnostic’ for sponges, the relevant question becomes: given the biosynthetic complexity and phylogenetic distribution of C-24 alkylation capacity, what is P(sponge | C_30_ sterane) in a given depositional context? This probabilistic framing accommodates the rhizarian finding without abandoning the compound’s utility as a sponge indicator in appropriate settings.

### Biosynthetic diversity reflects evolutionary history

The distribution of compounds in ψ– λ–σ space reflects evolutionary patterning of biosynthetic pathways recorded in MetaCyc. High retrobiosynthetic complexity indicates that many alternative routes exist to reach a product, assembled from experimentally characterized reactions that may occur in different lineages. This route diversity weakens source inference at the pathway level: detecting a sterol or sterane in sediment provides limited constraint on which biosynthetic route produced it, even before considering which organisms encode those routes.

Sterols illustrate how lineage-specific pathway evolution can generate distinct complexity signatures for closely related products. In the lanosterol-derived sterol network (animals and fungi), post-cyclization modifications support many alternative routes to downstream sterols, yielding high ψ and generally lower pathway consensus. In contrast, plant and algal sterol biosynthesis proceeding from cycloartenol is comparatively constrained, producing low-ψ sterols whose pathways share substantial structure. In this framework, differences among sterols reflect the evolutionary history of the pathway architecture itself, not just differences among target molecules. This interpretation is consistent with recent work showing that the Proterozoic sterane record reflects stepwise evolution of sterol biosynthesis and early biochemical complexity preceding later ecological expansion, rather than simple tracking of organismal abundance^20,32^.

Similar contrasts appear in tetrapyrrole pigments: a conserved core pathway is shared broadly^33,34^, while lineage-specific elaborations introduce additional branching and alternative routes for some pigments. These examples indicate that ψ, λ, and σ capture pathway diversification layered onto conserved biochemical cores

### Accumulation of complexity along pathways

Individual pathways show how complexity accumulates mechanistically along biosynthetic trajectories. In the lanosterol-derived sterol network, early cyclization products such as lanosterol show relatively low ψ, whereas downstream sterols exhibit progressively higher ψ as alternative post-cyclization modifications multiply. This increase reflects branching among demethylation, double-bond isomerization, and side-chain reduction steps. The result is a combinatorial expansion of routes to late-stage products such as cholesterol.

In contrast, the cycloartenol-derived sterol pathway (plants and algae) remains constrained across much of the trajectory, maintaining low ψ. Under this architecture, some downstream products can show high λ and high σ because alternative routes, where present, tend to diverge late and share most intermediates, but the primary driver of low ψ is the limited number of viable routes in the pathway as a whole. Thus, differences in ψ among sterols primarily reflect differences in pathway architecture between lanosterol- and cycloartenol-derived networks, while λ and σ describe where and how any remaining ambiguity is distributed along those routes.

This same pattern extends beyond sterols: conserved cores can persist for long evolutionary intervals, while lineage-specific additions create new branch points and alternative routes that measurably increase ψ and often reduce σ.

### Limitations and future directions

Our framework quantifies biosynthetic ambiguity – one of three sequential sources of interpretive uncertainty in biomarker analysis (Box 1, Figure 1). Complete evaluation requires consideration of: (1) phylogenetic ambiguity, arising from multiple organisms possessing the same biosynthetic capacity; (2) biosynthetic ambiguity, addressed here, arising from multiple pathways producing the same molecule; and (3) diagenetic ambiguity, arising from multiple precursors converging on shared preservation products. These uncertainties compose sequentially: a molecule with constrained biosynthesis may nonetheless be a poor biomarker if the pathway is distributed across diverse taxa or if the compound derives diagenetically from multiple precursors.

#### Biosynthetic limitations

MetaCyc pathway coverage reflects curated biochemical knowledge. Some pathways are not known, and some characterized pathways are not yet represented. The goal is not to estimate absolute biochemical uncertainty but to quantify inference limits given existing metabolic annotations, which are the same resources implicitly used in biomarker interpretation. We augmented the network with 27 supplementary reactions for geologically important biomarkers with characterized pathways absent from MetaCyc. However, some geologically important biomarkers lack completely characterized synthetic pathways, including dinosterol^35^ (dinoflagellates), crenarchaeol^36^ (Thaumarchaeota), dihydroxyarchaeol^37^ (archaea), ladderane lipids^38,39^ (anammox bacteria), alkenones^40^ (haptophytes), highly branched isoprenoids^41^ (diatoms), and cheilanthanes^42^. Uncharacterized pathways, particularly in uncultured lineages, may alter complexity estimates as biochemical knowledge expands.

The current framework treats all enzymatically plausible pathways as equally probable. In reality, metabolic flux is unevenly distributed. Incorporating flux data to weight pathway contributions would refine complexity estimates, and environment-specific metrics could account for the producers actually present in a given setting.

#### Diagenetic ambiguity

Examples of diagenetic convergence are well-documented^37^. For example, phytane derives from chlorophyll phytol, archaeal ether lipids, and tocopherols^38^; thermal degradation of hopanoids generates products identical to cheilanthanes^42^. Such convergence means a single preserved hydrocarbon may have originated from biosynthetically distinct precursors. No systematic database of sedimentary organic reactions currently exists analogous to MetaCyc. Constructing such a database would enable diagenetic complexity metrics analogous to those presented here. Until then, our biosynthetic metrics provide minimum ambiguity estimates, as diagenetic convergence can only increase interpretive uncertainty.

#### Evolutionary considerations

The metabolic network in MetaCyc reflects modern biochemistry. For Precambrian biomarkers, pathway diversity has expanded over Earth history while other pathways may have gone extinct – the organisms producing paleorenieratane^43,44^ remain unknown despite the compound’s presence in ancient rocks. Our metrics therefore represent upper bounds on ambiguity for ancient systems with simpler networks but miss extinct and uncharacterized pathways. Compounds appearing biosynthetically constrained today may have been more diagnostic when fewer alternatives existed; conversely, compounds from evolutionarily recent pathways (e.g., angiosperm terpenoids) would be absent from pre-Mesozoic systems. Time-resolved complexity could be reconstructed by pruning networks to ancient reactions and integrating with molecular clock approaches.

#### Broader implications

Rather than replacing existing biomarker approaches, this framework provides a quantitative basis for weighting molecular evidence, enabling explicit probabilistic interpretation of biomarker signals rather than categorical assignments. Formalizing biomarker interpretation as inference through a noisy channel, where phylogeny, biosynthesis, and diagenesis represent sequential stages of information loss, opens a path from qualitative assessment toward probabilistic inference.

## CONCLUSIONS

Retrobiosynthetic complexity (ψ), normalized branch depth (λ), and fraction shared (σ) provide complementary metrics for evaluating the diagnostic reliability of molecular biomarkers. Analysis of 10,467 metabolites from MetaCyc reveals that biosynthetic specificity – the combination of few pathways, distal divergence, and extensive shared biosynthesis – is rare: only 714 compounds (13% of multi-pathway metabolites) meet all three criteria. Lipid biomarkers span this specificity space heterogeneously, with each compound class occupying a distinct region defined by its pathway architecture rather than its preservation chemistry.

Because diagnostic quality and preservation potential are approximately independent, the principal constraint on molecular paleontology is the limited chemical diversity sampled by preservable compound classes. Biomarker interpretation therefore requires integrating multiple lines of evidence rather than providing definitive source assignments. The metrics introduced here quantify biosynthetic ambiguity, i.e. pathway-level constraints on diagnostic reliability. Translating these constraints into organism-level source attribution requires the additional step of mapping pathways to taxa, a problem whose tractability depends on the very pathway diversity that ψ, λ, and σ quantify. Compounds with constrained biosynthesis warrant detailed phylogenetic analysis; compounds with high pathway diversity are intrinsically ambiguous regardless of how producers are distributed.

## METHODS

### Network construction and filtering

The global metabolic network was constructed from all small-molecule metabolism reactions in the MetaCyc database using Pathway Tools version 29.5^45,46^. At the time of analysis, the database contained 3,270 metabolic pathways comprising curated enzymatic reactions across 20,553 small-molecule compounds with defined chemical structures spanning all domains of life. The MetaCyc network was augmented with 27 supplementary reactions capturing recently characterized pathways for hopanoid side-chain modifications (7 reactions)^47–50^, hopanoid A-ring methylation (2 reactions)^14,51^, archaeal tetraether and GMGT synthesis (4 reactions)^28,52,53^, hydroxyarchaeol synthesis (1 reaction)^54^, macrocyclic archaeol synthesis (1 reaction)^28,53^, methylated GMGT synthesis (1 reaction)^55^, OH-GDGT and methylated GDGT biosynthesis (3 reactions)^56^, aromatic carotenoid biosynthesis (3 reactions)^57–59^, C_30_ sterol pathways (2 reactions)^60^, coprostanol biosynthesis (1 reaction)^61^, tetrahymanol biosynthesis (1 reaction)^62^, and isoarborinol cyclization (1 reaction)^63^. The hydroxyarchaeol supplementary reaction connects to digeranylgeranylglycerophosphate (DGGGP), the biochemically verified substrate of the MA0127 hydratase^54^. The subsequent geranylgeranyl reductase step to produce fully saturated hydroxyarchaeol is implicit in the single-step representation. These additions enable complexity analysis for geologically important biomarkers not fully represented in MetaCyc.

Superpathways were excluded to avoid double-counting reactions already present in their constituent base pathways. Each reaction was represented as a directed hyperedge connecting substrate metabolites to product metabolites, with edge direction indicating biosynthetic direction (precursor → product). The resulting network comprised 10,467 unique metabolites connected by 10,148 unique reactions.

Of the 10,467 compounds in the network, 9,865 (94.2%) had identifiable biosynthetic pathways. Compounds with more than 10,000 distinct biosynthetic paths (n = 725) were excluded as representing central rather than specialized metabolism, e.g. tracing through glycolysis, the TCA cycle, and pentose phosphate pathway, yielding path counts orders of magnitude higher than the ≤48 paths observed for all target biomarkers. After excluding these central metabolism intermediates and 602 compounds lacking identifiable pathways, 9,140 compounds remained for analysis. Among these, 4,510 (49.3%) had single biosynthetic pathways (*ψ* = 0 bits by definition), while 4,630 (50.7%) had multiple pathways with calculable complexity.

### Mathematical formulation for retrobiosynthetic complexity

Retrobiosynthetic complexity (ψ) quantifies the entropy, in bits, over alternative biosynthetic pathways to a compound. It is defined as ψ = log_2_(Npaths), where Npaths is the number of unique biosynthetic routes connecting a compound to primary metabolic precursors. This metric addresses a central challenge in interpreting fossil hydrocarbons: when a molecule is detected in ancient rocks, its biological source must be inferred by reasoning backward from product to precursors. This retrobiosynthetic perspective reverses the conventional direction of metabolic pathway analysis. Many biomolecules lie at the convergence of multiple biosynthetic pathways, each involving distinct enzymatic reactions that may be distributed across different organisms. As the number of pathways leading to a molecule increases, so does the uncertainty associated with its source.

#### Derivation from decision tree entropy

Retrobiosynthetic complexity adapts the information-theoretic path entropy measure developed by Green^64^ for decision trees to metabolic hypergraphs. Green defined path entropy for decision trees, where each outward-directed path from the root to a leaf accumulates uncertainty by summing the base-2 logarithm of branching choices. For a tree with *n* complete paths:

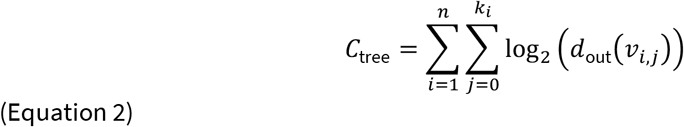

where *C*_tree_ is the total path complexity in bits, *v*_*i,j*_ denotes the *j*th node on the *i*th path, and *d*_out_(*v*_*i,j*_) is the out-degree of that node.

For metabolic networks, we reverse edge orientation^65,66^. Metabolic reactions proceed from precursors to products (edges directed forward), but in retrosynthesis we trace backward from the biomarker of interest (the root) to terminal compounds (the leaves). Reversing edge direction transforms the equation to:

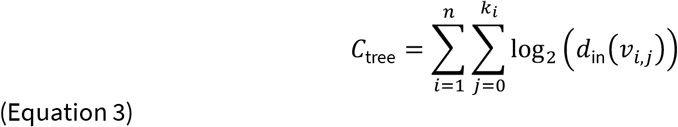

where *d*_in_(*v*_*i,j*_) is now the in-degree (number of edges pointing toward the node).

#### Extension to hypergraphs

The tree formulation captures uncertainty from multiple pathways but cannot represent convergent metabolic reactions where multiple substrates are required simultaneously^67^. When metabolite C is produced from A + B → C, and A can be synthesized through *m* pathways while B through *n* pathways, the total routes to C is *m* × *n*. This multiplicative relationship requires hypergraph representation^67,68^.

In a hypergraph, each reaction is a hyperedge connecting all substrates to all products, preserving the requirement that certain metabolites must be present simultaneously. The hypergraph-based retrobiosynthetic complexity is:

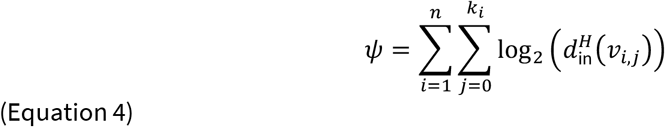

where *n* is the number of complete retrosynthetic paths, *k*_*i*_ is the length of path *i*, and 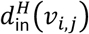 is the hypergraph in-degree – the number of reaction hyperedges that could produce metabolite *v*_*i,j*_.

#### Hub filtering and path termination

Currency metabolites participate in numerous reactions and would cause combinatorial explosion in path enumeration. We use connectivity-based filtering^65,69^: hub metabolites with out-degree > 20 are removed from reactant lists. This threshold, selected through systematic analysis across seven values (10–50), identified 198 hub metabolites including PROTON (out-degree = 7,405), WATER (4,697), ATP (2,767), and ACETYL-COA (2,297). β-carotene, despite having out-degree = 24, was protected from exclusion as an essential intermediate in carotenoid biosynthesis pathways.

Hub metabolites are removed from reactant lists but can still be analyzed as products. Biosynthetic paths terminate at the first hub metabolite encountered, measuring complexity from the first non-currency intermediate to the target compound.

#### Biochemically plausible versus experimentally confirmed pathways

The enumerated pathways represent biochemically plausible routes constructed from experimentally characterized reactions in MetaCyc^45^. Complete pathways may combine reactions from different organisms and may not represent naturally occurring routes in any single extant species. However, this diversity quantifies biosynthetic promiscuity across all known biology. Compounds accessible through many theoretical routes can be synthesized through diverse biochemical strategies; compounds with few routes have inherently constrained biosynthetic access. This pathway diversity determines the theoretical ambiguity in source attribution, independent of the phylogenetic distribution of biosynthetic capacity.

#### Interpretation

Retrobiosynthetic complexity, expressed in bits, quantifies cumulative decision complexity across all possible routes to a metabolite. The measure reflects pathway diversity rather than pathway length: a biomarker produced by a single long pathway has ψ = 0 (no ambiguity), while one synthesized through many short pathways has higher complexity. The approach uses uniform priors over experimentally characterized pathways. Compounds with low ψ provide reliable information about biological origins; those with high ψ require additional evidence for confident interpretation.

### Normalized branch depth formulation

Normalized branch depth quantifies the extent to which alternative biosynthetic routes share enzymatic machinery. While retrobiosynthetic complexity (*ψ*) measures pathway diversity, normalized branch depth (*λ*) measures where pathways diverge. Values range from 0 (pathways diverge proximally, i.e., immediately near the biomarker, indicating minimal shared biosynthesis) to 1 (pathways diverge distally, only near terminal metabolites, indicating extensive shared biosynthesis).

#### Branch point identification

For each metabolite *v* in a pathway, we define its branching contribution as:

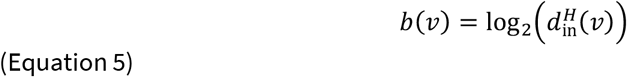

where 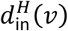 is the hypergraph in-degree (number of reactions producing *v*). Metabolites with 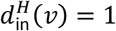 (single precursor) contribute zero to branching and represent pathway segments with no alternatives. Metabolites with 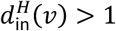 are branch points where pathway alternatives diverge.

#### Branch point depth

For each branch point *v*, we calculate its normalized depth from the biomarker:

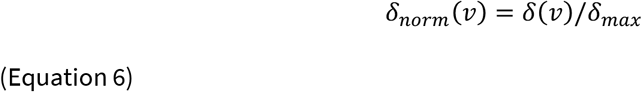

where *δ*(*v*) is the distance from the biomarker to *v* and *δ*_*max*_ is the maximum depth in the biosynthetic network. Values near 0 indicate positions close to the biomarker (minimal shared biosynthesis) and values near 1 indicate positions near terminal metabolites (extensive shared biosynthesis).

#### Normalized branch depth calculation

Normalized branch depth is the path-weighted mean of normalized branch point depths:

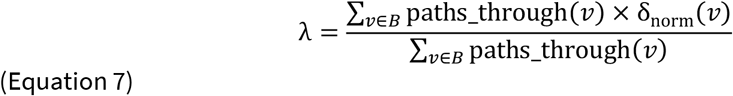

where *B* is the set of all branch points and paths_through(*v*) is the number of biosynthetic routes passing through branch point *v*. This weighting ensures that branch points traversed by many pathways contribute proportionally more to *λ* than branch points on rare routes. Normalized branch depth is normalized to [0,1]. High *λ* indicates pathways share extensive biosynthetic machinery (branch points far from biomarker), while low *λ* indicates minimal shared biosynthesis (branch points near biomarker).

For compounds with no branch points (single linear pathway), *λ* is undefined and these compounds are reported as having *ψ* = 0 rather than quantifying convergence.

### Fraction shared formulation

Fraction shared (σ) quantifies the extent to which alternative biosynthetic routes traverse common intermediates. While λ measures where pathways diverge, σ measures how much of each route uses consensus nodes – metabolites appearing in a high proportion of all pathways to a target compound. Crucially, σ depends on both branching position and pathway count: two compounds with similar λ can have very different σ if one has few paths (yielding high overlap among routes) while the other has many paths (diluting any single node’s representation across routes).

#### Consensus node identification

For each metabolite v in the biosynthetic network leading to a target compound, we calculate its frequency as the fraction of all pathways that pass through

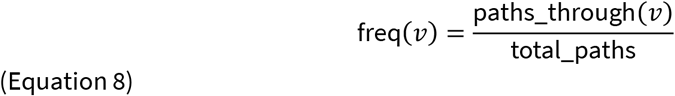

where paths_through(*v*) is the product of paths from terminal metabolites to *v* and paths from *v* to the target compound. A node is classified as a consensus node if freq(*v*) ≥ τ, where τ is the consensus threshold. We use τ = 0.80 as the default threshold, identifying nodes appearing in at least 80% of all biosynthetic routes.

#### Fraction shared calculation

The fraction shared metric is calculated as:

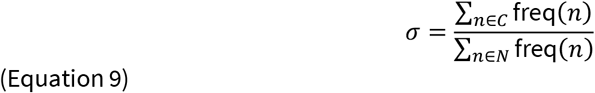

where C is the set of consensus nodes and N is the set of all nodes in the biosynthetic network. This gives the expected fraction of any pathway that uses consensus nodes.

#### Interpretation

The fraction shared metric partitions pathway structure into components that carry different amounts of information about biosynthetic origin. Consensus nodes (those appearing in ≥80% of pathways) represent shared biosynthetic machinery that provides no discrimination among alternative routes. Non-consensus nodes represent the divergent portions of the network where different biosynthetic routes are distinguishable.

High σ (> 0.5) indicates that pathways share a common biosynthetic backbone, concentrating whatever source-diagnostic information exists into the small fraction of the network where pathways diverge. Low σ (< 0.35) indicates that pathways proceed through largely independent sequences of intermediates. In information-theoretic terms, σ partitions the pathway into shared (source-agnostic) and unique (potentially diagnostic) components. The conditional entropy H(pathway | molecule) – residual uncertainty about biosynthetic route given the observed product – depends primarily on ψ, but σ determines where additional observations (e.g., pathway intermediates, isotopic signatures) would be most informative: for high-σ compounds, only the branch points carry discriminatory power.

### Dynamic programming algorithm

Retrobiosynthetic metrics were computed using a dynamic programming^70^ algorithm that calculates exact values in O(V+E) time without explicit path enumeration.

#### Main-reactant graph construction

For reactions with multiple reactants, only the first (main) reactant is considered for path traversal. The MetaCyc database consistently lists the primary biosynthetic substrate as the first reactant in reaction definitions, with cofactors (NADPH, ATP, etc.) listed subsequently. This ordering was validated for sterol, fatty acid, and terpenoid biosynthesis pathways: 100% of reactions for key biomarkers (cholesterol, ergosterol, squalene, palmitate, hopan-22-ol) had substrate-first ordering. This approach traces biosynthetic backbones without artificial path multiplication through cofactor biosynthesis.

This substrate-centric approach also ensures that reactions differing only in cofactor usage (e.g., NADPH versus NADH as electron donor, or NADPH versus ferredoxin) are counted as a single biosynthetic route, as they represent the same carbon skeleton transformation. For retrobiosynthetic complexity, the relevant question is how many distinct structural precursors can be converted into a product, not how many cofactor variants exist for the same transformation.

#### Hub filtering

The algorithm operates on the hub-filtered network described above, where metabolites with out-degree > 20 have been removed from reactant lists.

#### Cycle breaking

The MetaCyc reaction network contains cycles due to reversible reactions, metabolic cycles, and interconnected pathways. Cycles prevent topological sorting and cause infinite path counts. We detect and remove back-edges during depth-first search (DFS) traversal: edges to nodes already in the current recursion stack are identified as back-edges and excluded from the directed acyclic graph (DAG) used for DP computation.

Path counts are then computed on this DAG. For each node, we calculate the number of paths from terminal metabolites to that node (paths_to), the number of paths from that node to the target compound (paths_from), and the total number of paths that pass through the node, given by the product of these two quantities (paths_through = paths_to × paths_from). These values are computed via two DP passes: forward (from terminals toward target) and backward (from target toward terminals), each requiring O(V+E) time.

#### Metric computation

The three metrics are computed from path counts without explicit enumeration.

Retrobiosynthetic complexity:

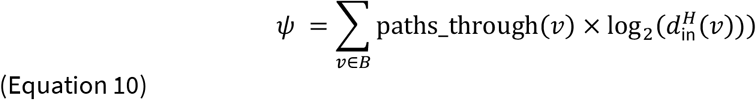

Where *B* is the set of branch points (nodes with in-degree > 1).

Normalized branch depth is computed as the path-weighted mean of normalized branch point depths (Equation 7), where *δ*_*norm*_(*v*) is the normalized depth of node *v* (distance from target divided by maximum depth). Fraction shared is computed as described above using node frequencies derived from paths_through values.

#### Computational performance

The algorithm successfully processed all 10,467 MetaCyc metabolites with a total runtime of ∼40 minutes (median 28 ms per compound). The linear time complexity enables analysis of the complete metabolic network without sampling or approximation.

### Statistical analysis

Complexity thresholds for biomarker classification were determined empirically from the distribution of retrobiosynthetic complexity values across all analyzed compounds. The distribution showed a natural break at ψ = 8 bits, separating low-complexity compounds (73.4% of the dataset) from high-complexity compounds (26.6%). Correlations between metrics were assessed using Pearson and Spearman coefficients. Independence was evaluated through decile analysis, examining how mean λ and σ varied across ψ deciles. Statistical analyses were performed using Python (NumPy, SciPy). Visualization was conducted using Matplotlib. All results were evaluated for robustness by examining sensitivity to the hub-filtering threshold across seven values (10–50).

## Acknowledgements

The authors acknowledge financial support from Washington University in the form of a graduate fellowship to M.H. They also thank David Fike, Paul Byrne, and Lon Chubiz for feedback that helped frame the study design.

## Author Contributions

Both authors contributed to all aspects of this work including conceptualization, methodology, software, analysis, and writing.

## Code Availability

Code used to calculate retrobiosynthetic complexity metrics and generate figures is available at https://github.com/bradleylab/retrobiosynthetic-complexity. The code will be archived in a public repository with a persistent identifier prior to publication and will be made available to editors and referees upon request during review.

## Data Availability

Retrobiosynthetic complexity values for all analyzed compounds are provided in Supplementary Data 1. The MetaCyc database is freely available at https://metacyc.org. Retrobiosynthetic complexity, normalized branch depth, and fraction shared values for all 9,140 compounds passing network filtering are provided in Supplementary Table 2.

---

**Table S1. Biosynthetic metrics for all biomarker compounds**

Retrobiosynthetic complexity (ψ), normalized branch depth (λ), and fraction shared (σ) for 86 biomarkers.

**Table S2. Biosynthetic metrics for all compounds**

Retrobiosynthetic complexity (ψ), normalized branch depth (λ), and fraction shared (σ) for all analyzed compounds.

